# Modeling The Effect of Forest Fire and Deforestation on Mid-Sized Wild Cat (Felids) Occupancy in SE Asian Tropical Rainforest

**DOI:** 10.1101/2021.05.18.444700

**Authors:** Andri Wibowo

## Abstract

Felids are mammal groups that also experiencing the effects of forest fire and deforestation rate. By using camera detection method, two felid species, *Prionailurus bengalensis* and *Pardofelis marmorata*, of tropical rainforests in SE Asia have been studied. The studied area was a rainforest in Sumatra that has experienced several forest fires with annual deforestation rates of 1.69%-2.89%. Occupancy model using Akaike Information Criterion (AIC) is in agreement that deforestation rate is the best explanatory covariate explaining the declining occupancy of those felid species. *P. marmorata* was known more sensitive to the both deforestation rate and forest fire frequency covariate effects since it has similar AIC values. While *P. bengalensis* was slightly affected by forest fires. Values of Area Under The Curve (AUC) of Receiver Operating Characteristic (ROC) were > 0.5 and these indicate adequate probability of forest fire effects on felid occupancy. Cut off value of occupancy of *P. bengalensis* was higher than *P. marmorata*. For *P. bengalensis*, the cut off value was 1.75 leading to a sensitivity and specificity of 62%. This is the threshold value for the prediction of numbers of *P. bengalensis* individual occurred where both sensitivity and specificity are maximized and as an effect of forest fire, and this can be used to classify areas as occupied by *P. bengalensis*.

## Introduction

Forest fire and deforestation rates are considered as two determinant factors driving animal assemblages mainly mammals in South East Asia region. In tropical forest in SE Asia, mammals also experience changes in post-fire habitat that affects the composition of the mammal community in the post-fire area. Forest fires combined with deforestation with large intensity and severe impacts can lead to poor habitat quality for some species that are unable to adapt and consequently reduce the diversity of species of mammal communities in ecosystems (Lyon et al. 2000, Fontaine & Kennedy 2012).

While the ecological impact of fires on forest ecosystems has been studied in various regions, limited information can be obtained regarding the impact of fires on tropical biodiversity. In Indonesia, forest fires combined with deforestations occur almost every year on a fluctuating scale on the land and forest areas. Considering as a mega biodiversity country, it is important to understand the impact of fire on mammals.

Regarding mammal’s biodiversity in Indonesia, several islands like Sumatra, Kalimantan, Sulawesi and Papua are known as the habitats of several endemic mammals. One of the important mammal groups are belong to felid mammals. In this family, one of the species is already known as flag species, which is the Sumatran tiger *Panthera tigris*. Beside this charismatic species, there are also several threatened felid species, for example leopard (*Panthera pardus*), sunda clouded leopard (*Neofelis diardi*), marbled cat (*Pardofelis marmorata*), golden cat (*Profelis aorata*), leopard cat (*Prionailurus bengalensis*), and asiatic golden cat *(Catopuma teminckii*). Despite there are growing contemporary researches on felids, however how those felid species responding to the forest fires and deforestations are still limited.

## Methodology

### Study area

The surveyed area was Bukit Barisan Selatan national park located in the bordered between Bengkulu and Lampung provinces with geocoordinates of 5°26'S 104°20'E. This park has a total area of 3,568 km^2^ (3568000 Ha) and stretching along the Bukit Barisan mountain range is in average only 45 km wide but 350 km long. The park’s northern part is mountainous landscape with its highest point while its southern section is a peninsula covered by montane forest, lowland tropical forest, coastal forest, and mangrove forest. The park was threatened by land use changes from tropical rainforest converted to cropland and the land use alteration was done by slash and burn method (Figure 1) leads to the forest fire occurrences. These threats were occurred in the boundary of the parks. From 2010 to 2014, the forest fire was absent. While there were 7 cases of forest fires in 2015. As a result, mainly the eastside of the parks in north and south have experienced forest cover loss (Figure 1). The deforestation rates were ranging from 1.69% to 2.89%.

**Figure 1.**
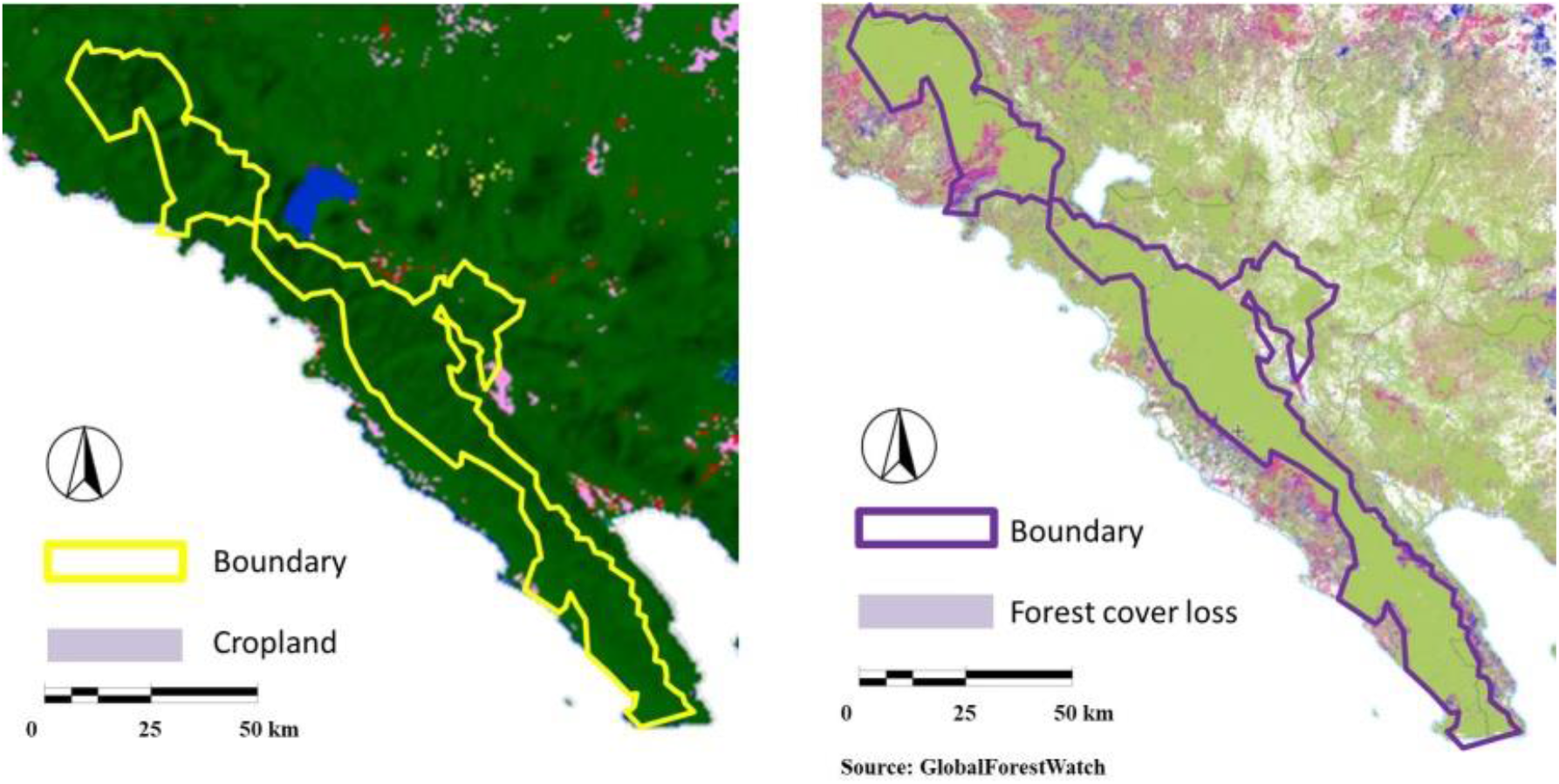
Boundary of Bukit Barisan Selatan NP bordered with cropland and forest cover loss

### Camera trap assembly

The presences of felids in study area were recorded by using camera traps (Figure 2) (Cheyne, 2012; Kuncahyo, 2017; Subagyo, 2019). The camera traps were installed at tree trunks at 30-40 cm height. To allow the identification of frontal and lateral body of species, the cameras were also positioned perpendicular to animal paths at 3-4 m distance. Likewise, the distance between one camera unit to another unit in grid cells was 1 km.

**Figure 2.**
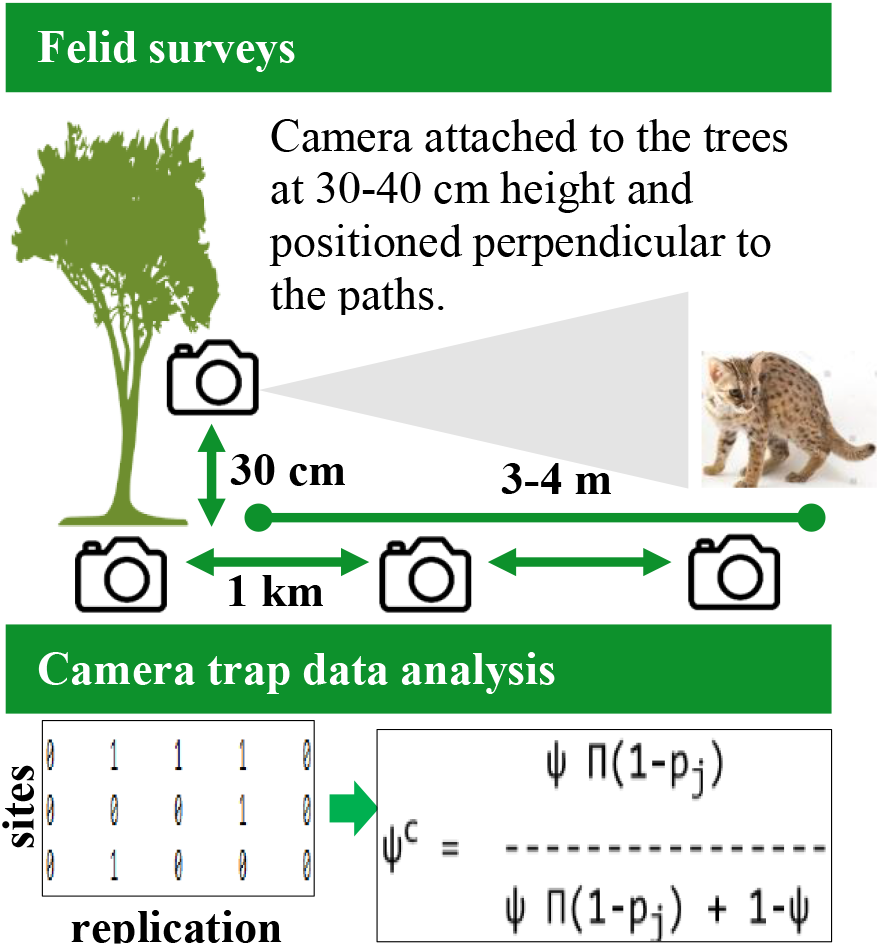
Camera trap assembly

### Camera data quantitative analysis

The quantitative analysis was performed based on the proportion of total sites occupied by the felid species denoted as occupancy or ψ (Wheeler and Dwiyahreni, 2007). The camera data in the form of identified felid pictures were extracted and tagged as 0 = the felid was absent/non-detection and 1 = the felid was present/a detection per replicated surveys in each camera trap site. The replicated survey histories for each species or species groups were constructed in a format with rows representing encounter histories at each site and columns representing detection at each sampling occasion. The occupancy of felids (ψ) was calculated based on this following Mackenzie’s equation:

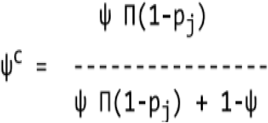

### Occupancy (ψ) and AIC modeling

Occupancy (ψ) modeling methods were based on Akaike Information Criterion (AIC) following Hellman (2013). Coleman et al. (2014), and Starbuck et al. (2015). AIC is simple to compute and easy to understand, and more importantly, for a given data set, it provides a measure of the strength of evidence for each model that represents a plausible biological hypothesis relative to the entire set of models considered. A feature of occupancy modeling is the ability to account for imperfect detection with the incorporation of detection covariates. The occupancy analysis was performed upon comparisons of events with presences of species and total sampling events. The occupancy variables were denoted as occupancy (ψ) and occupancy analyses were performed to compare amphibian occupancy in study area as functions of hydrosphere covariates.

To accomplish this, there is a two-stage approach. First, a series of candidate detection models were created to explain varying occupancies for felid species. The models ψ(.) contain an occupancy ψ(.) component were developed following work by MacKenzie et al. (2002). The models were tested using multi-model inference developed by Hines (2006). Then, a model set was proposed to explain occupancy of a species. The model set contained both a null and a global model incorporating covariates including forest fire frequency and deforestation rate. Forest fire frequency tests the impact of forest fire frequency on felid occupancy. Deforestation rate tests the impact of deforestation rate on felid occupancy.

Felid’s occupancy model as functions of forest fire frequency and deforestation rate was developed using AIC. The AIC was developed using the linear regression. The measured parameters included in AIC are ΔAIC and AIC weight. To build the model, 2 explanatory covariates including forest fire frequency, deforestation rate, and combinations of those covariates were included in the analysis to develop the model.

### Receiver Operating Characteristic (ROC)

Data used to analyze ROC were using measured variables consisting of felid occupancy, deforestation rate, and forest fire frequency. ROC analysis is an important test aiming for assessing the accuracy or discrimination performance of quantitative tests throughout the whole range of variables under experimental design (Calì and Longobardi, 2015). ROC analysis may also serve to estimate the accuracy of multivariate probability scores aimed at categorizing variables as affected/unaffected by a given treatment including deforestation rate and forest fire frequency. ROC was depicted as a curve and the results were measured based on the values of Area Under The Curve (AUC). The AUC is the ranking approach for assessing the performance, probability, and accuracy of the treatment on the measured variables. The performance of the treatment is demonstrated by the high value of the AUC, in which the AUC’s value of 0.5-0.7 is considered low, 0.7-0.9 as medium accuracy, and more than 0.9 indicates a high level of accuracy in measuring the treatment effect.

## Results & Discussion

Based on the conducted PCA analysis (Figure 3) between felid occupancy with deforestation rate and forest fire frequency covariates, certain negative associations were observed that those covariates have reduced felid species occupancy. *P. bengalensis* occupancy was higher than *P. marmorata*. Deforestation rate effect was more significant in comparison to forest fire. Three occupancy models tested for the *P. bengalensis* and *P. marmorata* species in Bukit Barisan Selatan national park can be seen in Table 1. The model is in agreement that deforestation rate is the best explanatory variable explaining the declining occupancy of those felid species. *P. marmorata* was known more sensitive to the both deforestation rate and forest fire frequency effects since it has similar AIC values. While *P. bengalensis* was slightly affected by forest fires.

**Figure 3.**
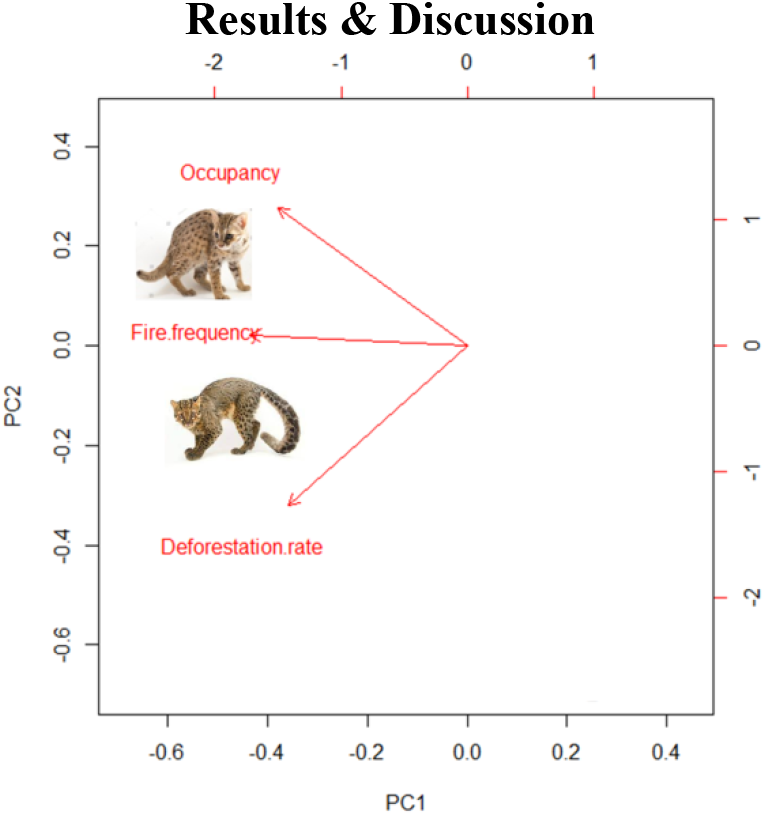
PCA of felid occupancy for *Prionailurus bengalensis* and *Pardofelis marmorata* with covariates of deforestation rate and forest fire frequency in Bukit Barisan Selatan national park.

**Table 1.** Three occupancy models tested for the *Prionailurus bengalensis* and *Pardofelis marmorata* with covariates of deforestation rate and forest fire frequency in Bukit Barisan Selatan national park (asterisk sign shows the best models).

| Felid occupancy ($\Psi$) + covariates | AIC | ΔAIC | AIC weight |
| --- | --- | --- | --- |
| *Prionailurus bengalensis* ($\Psi$) | | | |
| $\Psi$(fire frequency) | -13.50 | 1.73 | 0.29 |
| $\Psi$(deforestation rate) | -15.23* | 0.00 | 0.68 |
| $\Psi$(fire frequency + deforestation rate) | -9.23 | 6.00 | 0.03 |
| *Pardofelis marmorata* ($\Psi$) | | | |
| $\Psi$(fire frequency) | -10.73 | 0.01 | 0.47 |
| $\Psi$(deforestation rate) | -10.74* | 0.00 | 0.47 |
| $\Psi$(fire frequency + deforestation rate) | -6.45 | 4.28 | 0.06 |

Figure 4 and 5 show the Area Under The Curve (AUC) of Receiver Operating Characteristic (ROC) for estimating the probability of forest fire frequency effects on occupancy of both felid species. Both AUC values (Table 2) were > 0.5 and these indicate adequate probability of forest fire effects on felid occupancy. Cut off value of occupancy of *P. bengalensis* was higher than *P. marmorata*. For *P. bengalensis*, the plot of sensitivity and false positives (1-specificity) against expected probabilities indicates a probability cut off value of 1.75, leading to a sensitivity and specificity of 62%. This is the threshold value for the prediction of numbers of *P. bengalensis* individual occurred where both sensitivity and specificity are maximized and as an effect of forest fire, and this can be used to classify areas as occupied by *P. bengalensis* or not.

**Figure 4.**
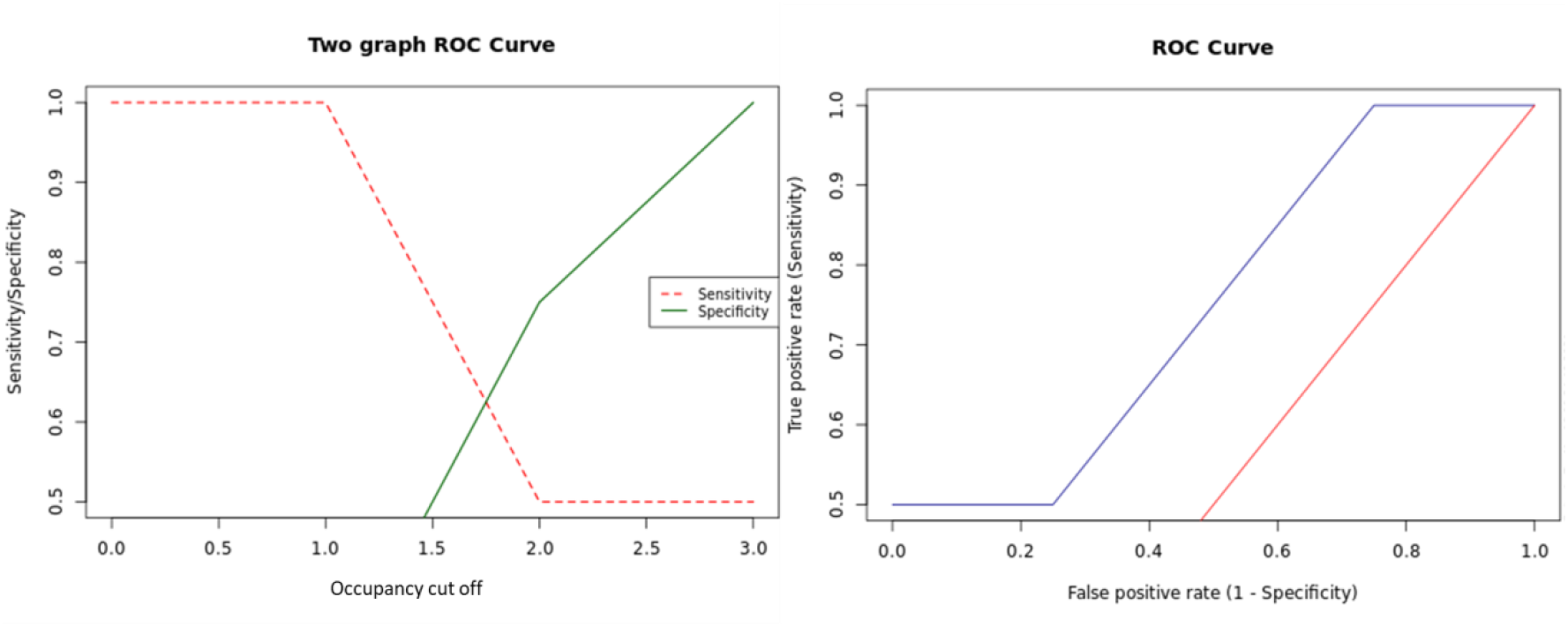
Area Under the Curve of ROC (AUROC) and occupancy cut off value representing the effects of forest fire frequency on *Prionailurus bengalensis* occupancy

**Figure 5.**
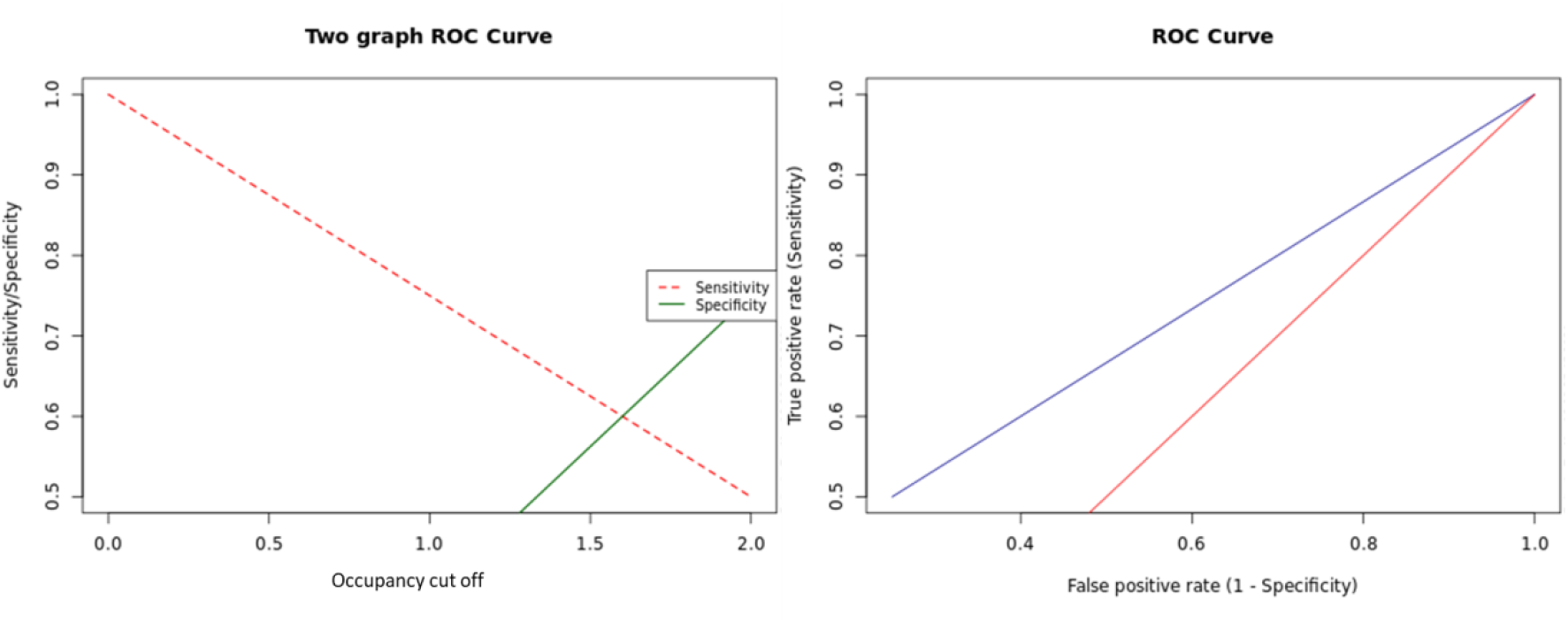
Area Under the Curve of ROC (AUROC) and occupancy cut off value representing the effects of forest fire frequency on *Pardofelis marmorata* occupancy

**Table 2.** Values of Area Under The Curve (AUC) of ROC (95% CI) for effect of forest fire frequency on *Prionailurus bengalensis* and *Pardofelis marmorata* occupancy

| Species | AUC | 95% CI |
| --- | --- | --- |
| *Prionailurus bengalensis* | 0.75 | 0.221-1 |
| *Pardofelis marmorata* | 0.625 | 0.077-1 |

## Notes

### Competing Interest Statement

The authors have declared no competing interest.

## References

Calì C, Longobardi, M. 2015. Some mathematical properties of the ROC curve and their applications. Ricerche mat.

Cheyne S, Ripoll B, Macdonald E, Sastramidjaja WJ. 2012. Standard operating procedure (sop) to install camera trap.

Coleman L, Ford W, Dobony C, Britzke E. 2014. Comparison of Radio-Telemetric Home-Range Analysis and Acoustic Detection for Little Brown Bat Habitat Evaluation. Northeastern Naturalist. 21. 431–445.

Deere, NJ, Guillera-Arroita, G, Platts, PJ, Mitchell, SL, Baking, EL, Bernard, H, Haysom, JK, Reynolds, G, Seaman D J I, Davies ZG, Struebig MJ. 2020. Implications of zero-deforestation commitments: Forest quality and hunting pressure limit mammal persistence in. fragmented tropical. Conservation Letters.

Fontaine J, Kennedy P. 2012. Meta-analysis of avian and small-mammal response to fire severity and fire surrogate treatments in U.S. Fire-prone forests. Ecological applications: a publication of the Ecological Society of America. 22: 1547–61.

Hellman ML. 2013. Amphibian Occupancy and Functional Connectivity of Restored Wetlands in the Missouri River Floodplain. Dissertations & Theses in Natural Resources. University of Nebraska-Lincoln. 68.

Hines JE. 2006. Presence 3.1-Software To Estimate Patch Occupancy And Related Parameters. USGS-PWRC.

Kuncahyo, BA, Alikodra, HA, Gunawan, H. 2017. identifikasi faktor sebaran macan dahan (neofelis diardi cuvier, 1823) di ekosistem rawa gambut, taman nasional Sebangau

Lyon L, Huff M, Hooper R, Telfer E, Schreiner D, Smith J. 2000. Wildland Fire in Ecosystems Effects of Fire on Fauna.

MacKenzie DI, Nichols JD, Lachman GB, Droege S, Royle JA, Langtimm CA. 2002. Estimating site occupancy rates when detection probabilities are less than one. Ecology 83:2248–2255.

Subagyo, A. Muhammad Y., Sumianto, Supriatna, J., Andayani, N., Mardiastuti, A., Sjahfirdi, L., Yasman, Sunarto. 2019. Survei dan monitoring kucing liar (carnivora:felidae) di taman nasional Way Kambas, Lampung, Indonesia.

Starbuck CA, Amelon SK, Thompson F. 2015. Relationships Between Bat Occupancy and Habitat and Landscape Structure Along a Savanna, Woodland, Forest Gradient in the Missouri Ozarks. Wildlife Society Bulletin 39(1).

Tilker A. 2014. Estimating Site Occupancy for Four Threatened Mammals in Southeastern Laos

Wheeler, P and Dwiyahreni, A. 2007. Large mammal monitoring in Lambusango.

